# Functional studies of deafness-associated pendrin and prestin variants

**DOI:** 10.1101/2024.01.23.576877

**Authors:** Satoe Takahashi, Takashi Kojima, Koichiro Wasano, Kazuaki Homma

## Abstract

Pendrin and prestin are evolutionary conserved membrane proteins that are essential for normal hearing. Pendrin is an anion transporter required for normal development and maintenance of ion homeostasis in the inner ear, while prestin is a voltage-dependent motor responsible for cochlear amplification essential for high sensitivity and frequency selectivity of mammalian hearing. Dysfunction of these proteins result in hearing loss in humans, and numerous deafness-associated pendrin and prestin variants have been identified in patients. However, the pathogenic impacts of many of these variants are ambiguous. Here we report results from our ongoing efforts in experimentally characterizing pendrin and prestin variants using *in vitro* functional assays, providing invaluable information regarding their pathogenicity.

## Introduction

The SLC26 family of membrane proteins includes two members, pendrin (SLC26A4) and prestin (SLC26A5), that are essential for normal hearing. Pendrin is an anion transporter required for normal development and maintenance of ion homeostasis in the inner ear (Choi et al., 2011; Everett et al., 2001). Prestin is a voltage-dependent motor and mediates voltage-dependent somatic motility (electromotility) of cochlear outer hair cells (OHCs), which is essential for high sensitivity and frequency selectivity of mammalian hearing (Cheatham et al., 2004; Dallos et al., 2008; Liberman et al., 2002; Zheng et al., 2000). Dysfunction or loss of these SLC26 proteins result in hearing loss. Pendrin-coding *SLC26A4* is a causative gene for Pendred syndrome (PDS, OMIM 274600), which is one of the most common types of syndromic hearing loss with enlarged vestibular aqueduct (EVA) and goiter, as well as nonsyndromic hearing loss (DNFB4, OMIM600791) (Everett et al., 1997; Pryor et al., 2005). Prestin-coding *SLC26A5* is associated with nonsyndromic hearing loss (DNFB61, OMIM613865). Since the advent of next-generation sequencing, large numbers of deafness-associated variants have been identified in patients. As of January 2024, 704 and 16 variants have been identified in *SLC26A4* and *SLC26A5*, respectively (The Human Gene Mutation Database, (Stenson et al., 2020)). However, experimental assessments on functional phenotypes of these genetic variants have been lagging behind to determine their pathogenicity with confidence. Previously, we have quantitated the anion transport functions of disease-associated pendrin variants in order to understand the pathology underlying various severity of PDS and DFNB4. (Wasano et al., 2020). Similarly, we characterized the functional phenotypes of prestin variants in both *in vitro* and *in vivo* to determine their pathogenicity (Takahashi et al., 2016; Takahashi et al., 2023a). Those experimental efforts are indispensable to provide information regarding effects of variants on protein functions, which in turn improves consistency and accuracy of genetic diagnoses of patients. This is in line with the American College of Medical Genetics and Genomics (ACMG) and the Association for Molecular Pathology (AMP) guidelines for the interpretation of sequence variants, adapted for genetic hearing loss (Oza et al., 2018).

In this study, we continue to assess the pathogenicity of deafness-associated pendrin and prestin variants *in vitro* to determine their roles in hearing loss in patients. For pendrin, highly reproducible fluorometric antiport assays previously developed by our group were used to interrogate the functional impacts of pendrin variants on the anion transport function (Wasano et al., 2020), with a major focus on transmembrane (TM) 9 and 10 located within the core domain that accommodates an anion substrate and undergoes elevator-like motions with respect to the gate domain during the transport cycle. For prestin, nonlinear capacitance (NLC) measurement was used to determine the functional impacts of a few prestin variants reported to date on the motile function (Ashmore, 1990; Santos-Sacchi, 1991). In addition, we compared our experimental results with predictions from AlphaMissense (AM) (Cheng et al., 2023), a recently developed *in silico* tool that predicts variant effects using artificial intelligence. We found decent correlation between AM pathogenicity scores and the antiport activity of pendrin variants, indicating good overall performance of this prediction tool (ρ = -0.6497). However, some of the variants were mis-categorized, indicating that prediction tools still have limitations and the experimental efforts remain important to assess the pathogenicity of deafness-associated variants.

## Methods

### Generation of stable cell lines

cDNAs coding human pendrin and gerbil prestin, (WT and variants) with a mTurquoise (mTq2) tag on the C-terminus were cloned into a pSBtet-pur vector (Addgene) using SfiI sites. Stable cell lines carrying these constructs were established in HEK293T cells as previously described (Wasano et al., 2020). Stable cells were maintained in DMEM supplemented with 10% FBS and 1µg/ml puromycin (Fisher Scientific), and the expression of the pendrin and prestin constructs were induced by the addition of doxycycline (Dox) in the media. For I^−^/Cl^−^ antiport assay, the pendrin-expressing pSBtet-Pur vectors were transfected in the HEK293T cell line that constitutively express mVenus^p.H148Q/p.I152L^ as previously described (Wasano et al., 2020).

### Fluorometric anion transport assays

Fluorometric HCO_3_^-^/Cl^-^ and I^-^/Cl^-^ antiport assays were established and described in detail in a previous study (Wasano et al., 2020). Briefly, for HCO^3-^/Cl^-^ antiport assay, stable HEK293T cells expressing pendrin constructs were loaded with a pH indicator, SNARF-5F (S23923, Thermo Fisher Scientific) in a high chloride buffer containing (mM): 140 NaCl, 4.5 KCl, 1 MgCl_2_, 2.5 CaCl_2_, 20 HEPES (pH 7.4). The antiport assay was initiated by an automated injection of a low chloride buffer containing (mM): 125 Na-gluconate, 5 K-gluconate, 1 MgCl_2_, 1 CaCl_2_, 20 HEPES, 25 NaHCO_3_ with 5% CO_2_ in Synergy Neo2 (Agilent/BioTek). The fluorescence of SNARF-5F were measured in a time dependent manner using Synergy Neo2 (Agilent/BioTek) and the data analyzed offline as described previously (Wasano et al., 2020). For I^-^/Cl^-^ antiport assay, cells expressing both pendrin variants and iodide sensitive fluorescent protein, mVenus^p.H148Q/p.I152L^, were resuspended in 200 μl of a high Cl^−^ buffer containing (mM): 150 NaCl, 1 MgCl_2_, 1 CaCl_2_, 20 HEPES, (pH 7.5). The I^−^/Cl^−^ antiport assay using 160 µl of the cell suspension was initiated by an automated injection of 80 μl of a high I^−^ buffer containing (mM): 150 NaI, 1 MgCl_2_, 1 CaCl_2_, 20 HEPES (pH 7.5) in Synergy Neo2 (Agilent/BioTek). The fluorescence intensities of mVenus^p.H148Q/p.I152L^ and mTq2 were simultaneously measured in a time-dependent manner using Synergy Neo2 (Agilent/BioTek) and the data analyzed offline as described previously (Wasano et al., 2020).

### Whole-cell recordings

Whole-cell recordings were performed at room temperature using the Axopatch 200B amplifier (Molecular Devices) with a 10 kHz low-pass filter. Recording pipettes pulled from borosilicate glass were filled with an ionic blocking intracellular solution containing (mM): 140 CsCl, 2 MgCl_2_, 10 EGTA, and 10 HEPES (pH 7.4). Cells were bathed in an extracellular solution containing (mM): 120 NaCl, 20 TEA-Cl, 2 CoCl_2_, 2 MgCl_2_, 10 HEPES (pH 7.4). Osmolality was adjusted to 309 mOsmol/kg with glucose. Holding potentials were set to 0 mV. NLC was measured using sinusoidal voltage stimuli (2.5-Hz, 120 or 150 mV amplitude) superimposed with two higher frequency stimuli (390.6 and 781.2 Hz, 10 mV amplitude). Data were collected by jClamp (SciSoft Company, New Haven, CT) (Santos-Sacchi et al., 1998).

### NLC data analysis

Voltage-dependent C_m_ data were analyzed using the following two-state Boltzmann equation:

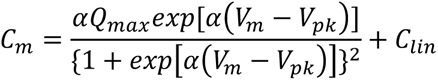

where α is the slope factor of the voltage-dependence of charge transfer, Q_max_ is the maximum charge transfer, V_m_ is the membrane potential, V_pk_ is the voltage at which the maximum charge movement is attained, and C_lin_ is the linear capacitance. The specific capacitance, C_sp_, was calculated as (C_m_ – C_lin_)/C_lin_.

### Statistical analyses

Statistical analyses for fluorometric antiport assays were performed as previously described (Wasano et al., 2020). Briefly, five different Dox dosage conditions were compared to the rates of uninduced cells by one-way ANOVA followed by uncorrected Fisher’s Least Significant Difference (LSD). To assess the dependency of the transport rates on Dox dosage, linear regressions (log_10_ [Dox] vs. transport rates) were performed. *F* tests were performed to find the difference in the Dox-dependence between WT versus pendrin variants. To obtain normalized activities of each pendrin variant, slope value from the linear regressions described above was divided by that of WT. The uncertainties (σ) associated with division computations to were calculated as previously described (Takahashi et al., 2023b) using the following equation:

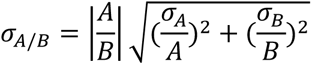

where A and B are the mean values with associated errors in linear regressions, σ_A_ and σ_B_, respectively. These values are reported as errors in **Table 2** under “%WT Activity”. The Spearman Correlation coefficient (ρ) between pendrin HCO_3_^-^/Cl^-^ transport activities and the AM scores was calculated using Prism. For NLC data analyses, one-way ANOVA combined with the Tukey’s post hoc test was used for multiple comparisons. In all statistical tests, *p* < 0.05 was considered statistically significant.

## Results

### Functional characterization of pendrin variants

A recent cryo-EM study on mouse pendrin (Liu et al., 2023) revealed a homodimeric architecture that is similar to prestin (Bavi et al., 2021; Butan et al., 2022; Futamata et al., 2022; Ge et al., 2021) and other SLC26 proteins (Chi et al., 2020; Tippett et al., 2023; Walter et al., 2019). The transmembrane domain consists of the gate and the core domains, and the ion translocation pathway is located in between them (**Fig. 1A**). The positive helix dipoles of TM3 and TM10 (their N-termini), together with a conserved basic residue (Arg^409^ and Arg^399^ in pendrin and prestin, respectively) in TM10 constitutes a positively charged pocket that holds a variety of anion substrates. The N-terminus of TM10 is connected to TM9 that is located on the opposite side of the core domain. The importance of the conserved residues in TM9, TM10, and the linker portion connecting these two helices (**Fig. 1B**) is suggested by the presence of multiple clustered missense variants identified in these regions: p.D380N (c.1138G>A), p.N382K (c.1146C>G), p.Q383E (c.1147C>G), p.Q383P (c.1148A>C), p.E384Q (c.1150G>C), p.E384G (c.1151A>G), p.I386V (c.1156A>G), p.A387V (c.1160C>T), p.G389R (c.1165G>A), p.G389R (c.1165G>C), p.S391R (c.1173C>A), p.S391N (c.1172G>A), p.N392S (c.1175A>G), p.N392Y (c.1174A>T), p.G396E (c.1187G>A), p.S399P (c.1195T>C), p.V402L (c.1204G>T), p.V402M (c.1204G>A), p.T404N (c.1211C>A), p.T404I (c.1211C>T), p.A406T (c.1216G>A), p.R409C (c.1225C>T), p.R409H (c.1226G>A), p.R409L (c.1226G>T), p.R409P (c.1226G>C), p.T410K (c.1229C>A), p.T410M (c.1229C>T), p.A411P (c.1231G>C), p.A411T (c.1231G>A), p.V412I (c.1234G>A), p.Q413R (c.1238A>G), p.Q413P (c.1238A>C), p.E414K (c.1240G>A), p.S415R (c.1245C>A), p.S415G (c.1243A>G), p.T416P (c.1246A>C) (Albert et al., 2006; Blons et al., 2004; Carvalho et al., 2018; Chai et al., 2013; Chen et al., 2007; Choi et al., 2009; Courtmans et al., 2007; Coyle et al., 1998; Elsayed and Al-Shamsi, 2022; Gonzalez Trevino et al., 2001; Huang et al., 2018; Hutchin et al., 2005; Ji et al., 2009; Koohiyan et al., 2021; Ladsous et al., 2014; Landa et al., 2013; Miyagawa et al., 2014; Park et al., 2003; Pera et al., 2008b; Reardon et al., 2000; Rendtorff et al., 2013; Sakuma et al., 2016; Sloan-Heggen et al., 2016; Turner et al., 2019; Van Hauwe et al., 1998; Wang et al., 2007; Wasano et al., 2020; Wen et al., 2019; Wu et al., 2005; Yuan et al., 2009; Zhao et al., 2014). Among them, p.E384G, p.N392Y, p.S399P, p.R409H, p.T410K, p.T410M, p.Q413R, p.Q413P, and p.T416P were functionally characterized and found to significantly impair the anion transport function of pendrin in previous studies by us and others (Choi et al., 2009; de Moraes et al., 2016; Gillam et al., 2005; Ishihara et al., 2010; Pera et al., 2008a; Scott et al., 2000; Taylor et al., 2002; Wasano et al., 2020; Yoon et al., 2008). The present study extended our experimental efforts to define the functional consequences of missense variants identified in the TM9-TM10 region. To this end, we established HEK293T stable cell lines expressing pendrin variants in a doxycycline (Dox) dosage-dependent manner and conducted HCO_3_^-^/Cl^-^ and I^-^/Cl^-^ antiport assays alongside wild-type (WT) control as described in detail in our previous report (Wasano et al., 2020). **Figs. 2A** and **2B** show the results of 25 pendrin missense variants evaluated here for their Dox dosage-dependent HCO_3_^-^/Cl^-^ and I^-^/Cl^-^ antiport activities. Many of the variants exhibited vastly reduced or no activity, with only a few had the WT-like activity (summarized in **Table 1**). These observations further affirm the importance of the conserved residues in TM9-TM10 in the core domain for the anion transport function of pendrin.

**Fig. 1.**
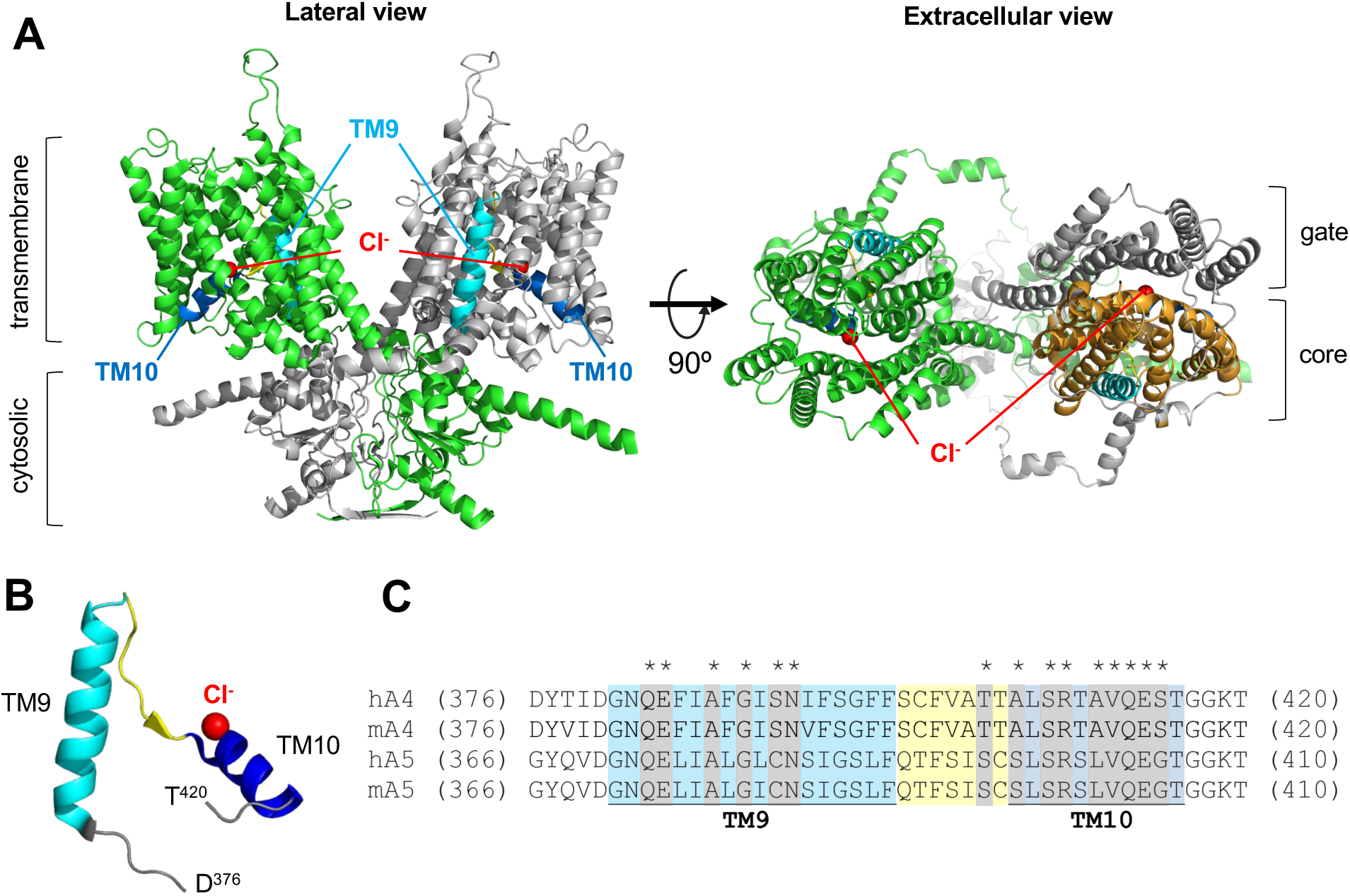
TM9-10 of pendrin. (**A**) The homodimeric structure of mouse pendrin (PDB: 7WK1). Protomers are shown in green and gray. Transmembrane and cytosolic domains are indicated in the lateral view (left). TM9 and TM10 are highlighted in cyan and blue, respectively, with connecting residues highlighted in yellow. The bound chlorides are indicated by red spheres. In the extracellular view (right), the core domain of one of the protomers is shown in bright orange. (**B**) TM9 and TM10 region of the structure (residues 376-420), extracted from the right protomer in (**A**). TM9 (381-398) is shown in cyan, linker region (399-405) is in yellow, and TM10 (406-416) is in blue. Bound chloride is shown as a red sphere. (**C**) Partial amino acid sequences of human and mouse pendrin (A4) and prestin (A5) showing the TM9 and TM10 region. Numbers in parentheses indicate the residue numbers at the N- and C-terminal ends of the partial amino acid sequences. TM9 and TM10 are highlighted in cyan and blue, respectively, with connecting residues highlighted in yellow as in (**A**). The residues with asterisks on top and gray shades indicate the locations of missense changes evaluated in Fig. 2.

**Fig. 2.**
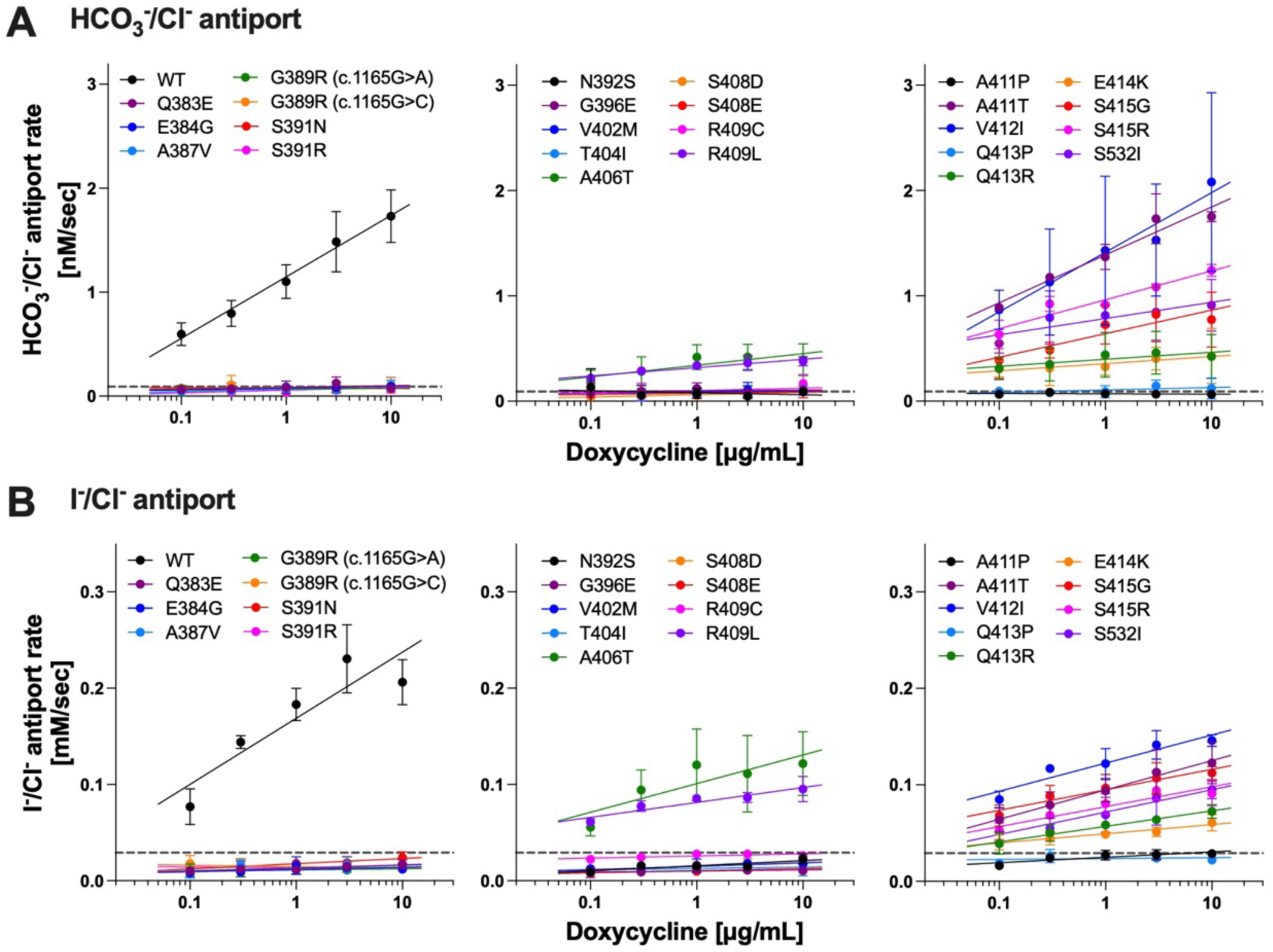
HCO_3_^-^/Cl^-^ and I^-^/Cl^-^ antiport assays on pendrin variants. HCO_3_^-^/Cl^-^ (**A**) and I^-^/Cl^-^ (**B**) antiport rates were plotted against doxycycline (Dox) concentration (0.1-10 µg/ml) for each mTq2-tagged pendrin variant alongside WT as indicated. Horizontal dotted lines indicate transport rates of uninduced cells. Error bars indicate SD. Solid lines indicate linear regressions (log_10_ [Dox] vs. transport rates). Sample size information and statistics are provided in Table 1.

**Table 1.**
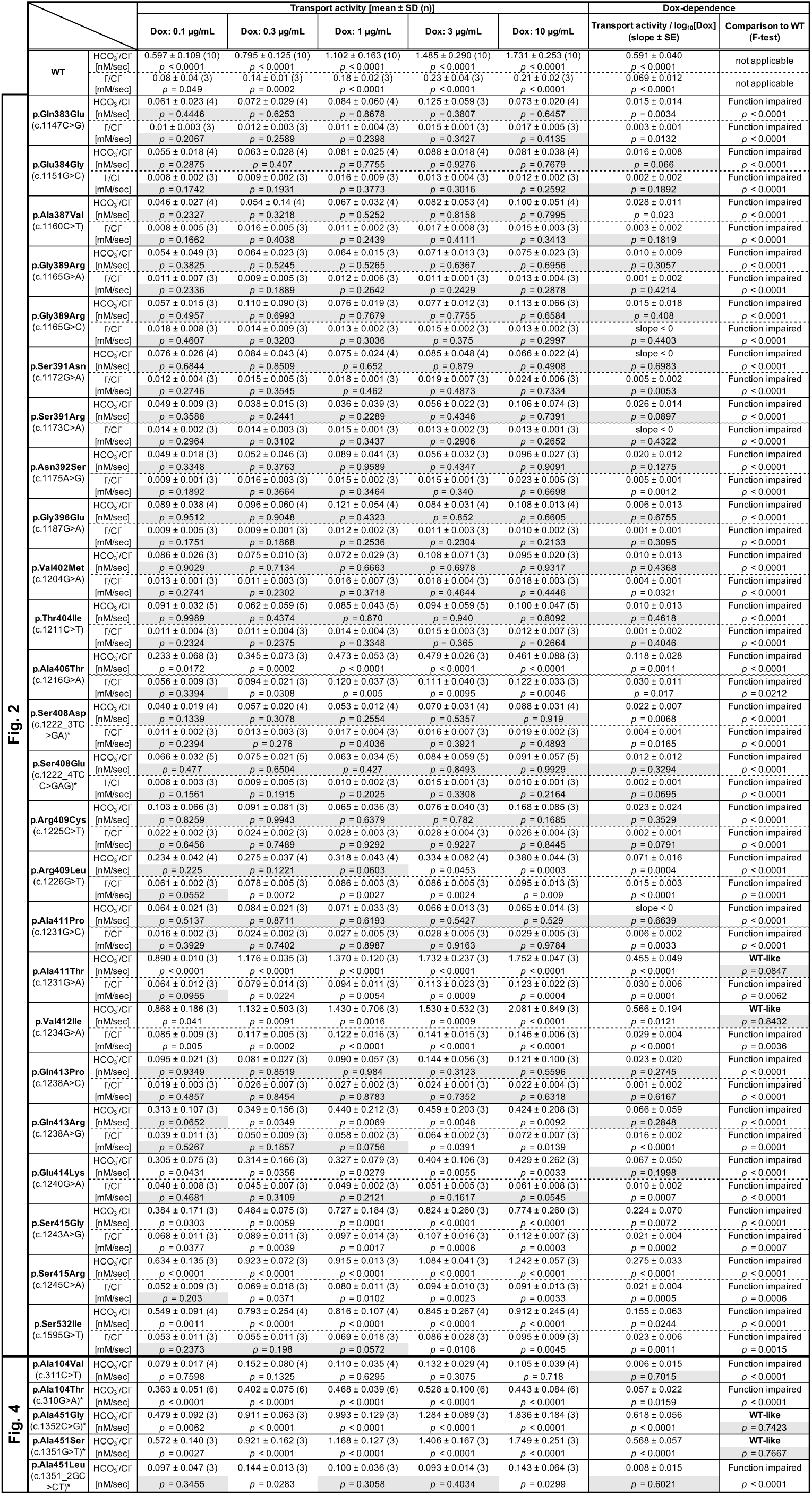
Summaries of the antiport rates of pendrin variants determined in this study. Numerical data from HCO_3_^-^/Cl^-^ and I^-^/Cl^-^ antiport assays in Fig. 2 and for HCO_3_^-^/Cl^-^ antiport assays in Fig. 4 are listed as indicated. *P*-values from one-way ANOVA with Fisher’s LSD tests compared to uninduced basal rates were listed below the mean ± SD values for each Dox concentrations. The numbers in the parentheses indicate the sample size. Comparison to WT was performed using the F-test on slope values as in our previous study (Wasano et al., 2020). Box with gray shades indicate statistically not significant (*p* ≥ 0.05). Asterisks (*) indicate variants not found in human patients (as of January 2024).

### Functional characterization of prestin variants

Mammalian prestin is unique among the SLC26 family of proteins, as it functions as a voltage-dependent motor instead of a transporter. Prestin also shares the overall similarities in the structure as pendrin and other SLC26 proteins, implying a common molecular principle underlying transport and motor functions. Thus, it is conceivable that conserved residues between pendrin and prestin are of similar functional importance in these structurally similar proteins. Interestingly, however, the number of prestin missense variants reported to date is only 13, of which 6 are associated with autism with unknown pathophysiological relevance to hearing (Cardenas et al., 2023; Han et al., 2019; Koire et al., 2021; Morgan et al., 2018; Mutai et al., 2013; Sloan-Heggen et al., 2016; Toth et al., 2007; Zhou et al., 2022). In this study, we characterized p.A100T (c.298G>A), p.P119S (c.355C>T), and p.S441L (c.1322C>T) prestin variants that are all associated with hearing loss. The locations of these three prestin missense variants are indicated in **Fig. 3A**. We established HEK293T cell lines expressing these prestin variants and measured the NLC as a proxy for their motor function. **Fig. 4A** shows the NLC of p.A100T and p.P119S prestin compared to WT. Although reduced, the NLC was detected for both p.A100T and p.P119S prestin, indicating some residual activity for these variants (**Fig. 4B**). These Ala and Pro residues are relatively well conserved among the SLC26 proteins (Ala^104^ and Pro^123^, respectively, in pendrin, **Fig. 3B**), and previous studies by us and others have shown that p.P123S variant significantly impair the anion transport function of pendrin (Ishihara et al., 2010; Wasano et al., 2020). In this study, we also measured the HCO_3_^-^/Cl^-^ antiporter activity for p.A104V pendrin (deafness-associated) and p.A104T pendrin (equivalent to p.A100T prestin but not identified in patients as of January 2024) and found that these two pendrin missense variants also significantly impair the anion transport function (**Fig. 5, left**), suggesting the common importance of these conserved residues for pendrin and prestin.

**Fig. 3.**
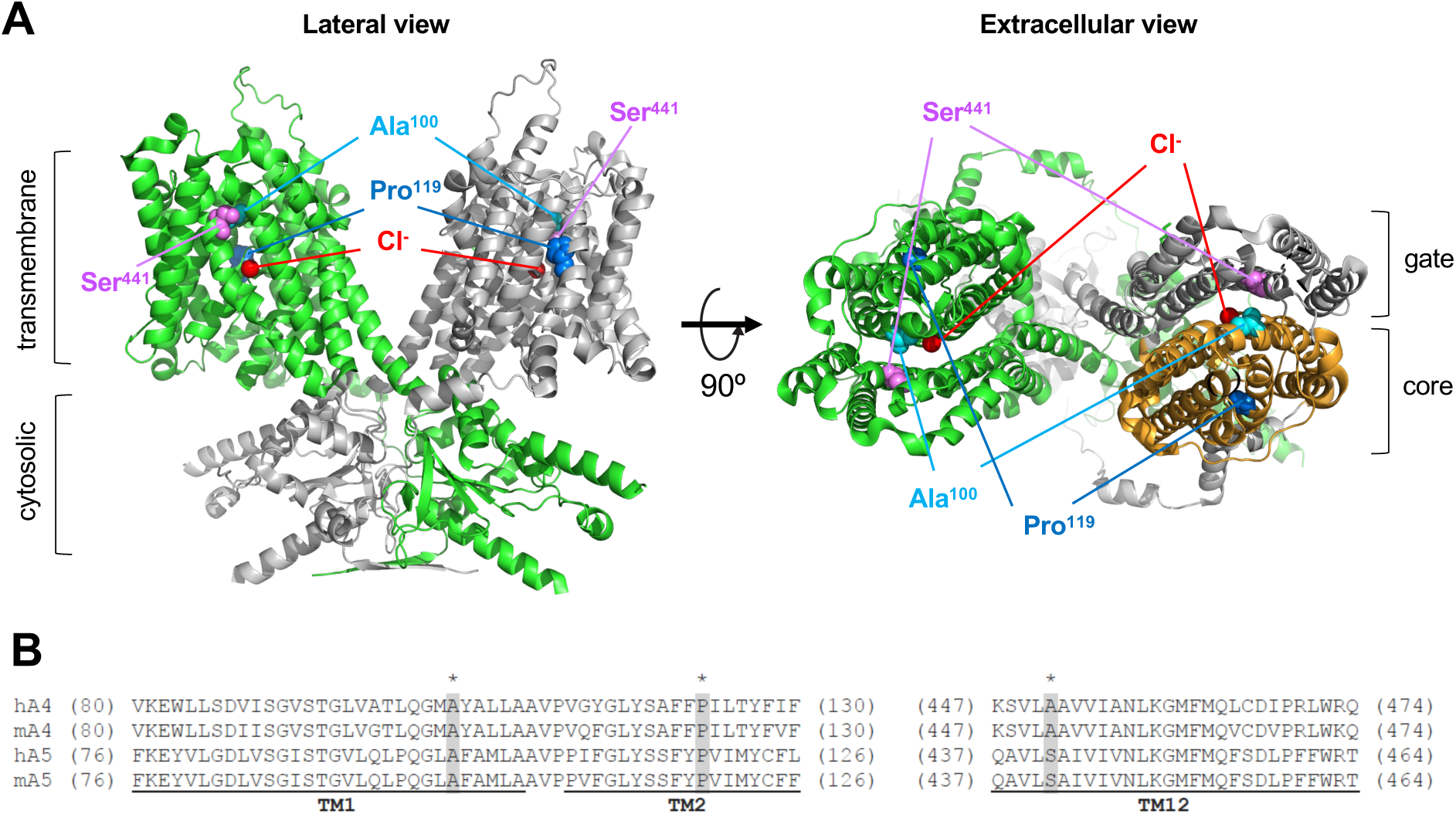
Ala^100^, Pro^119^, and Ser^441^ sites in prestin. (**A**) The homodimeric structure of human prestin (PDB: 7LGU). Protomers are shown in green and gray. In right panel, the core domain of one of the protomers is shown in bright orange. The Ala^100^, Pro^119^, and Ser^441^ sites and bound chlorides are indicated by cyan, blue, purple, and red spheres, respectively. (**B**) Partial amino acid sequences of human and mouse pendrin (A4) and prestin (A5). Numbers in parentheses indicate the residue numbers at the N- and C-terminal ends. The residues with asterisks indicate Ala^100^, Pro^119^, and Ser^441^ in human prestin (hA5) and equivalents in others.

**Fig. 4.**
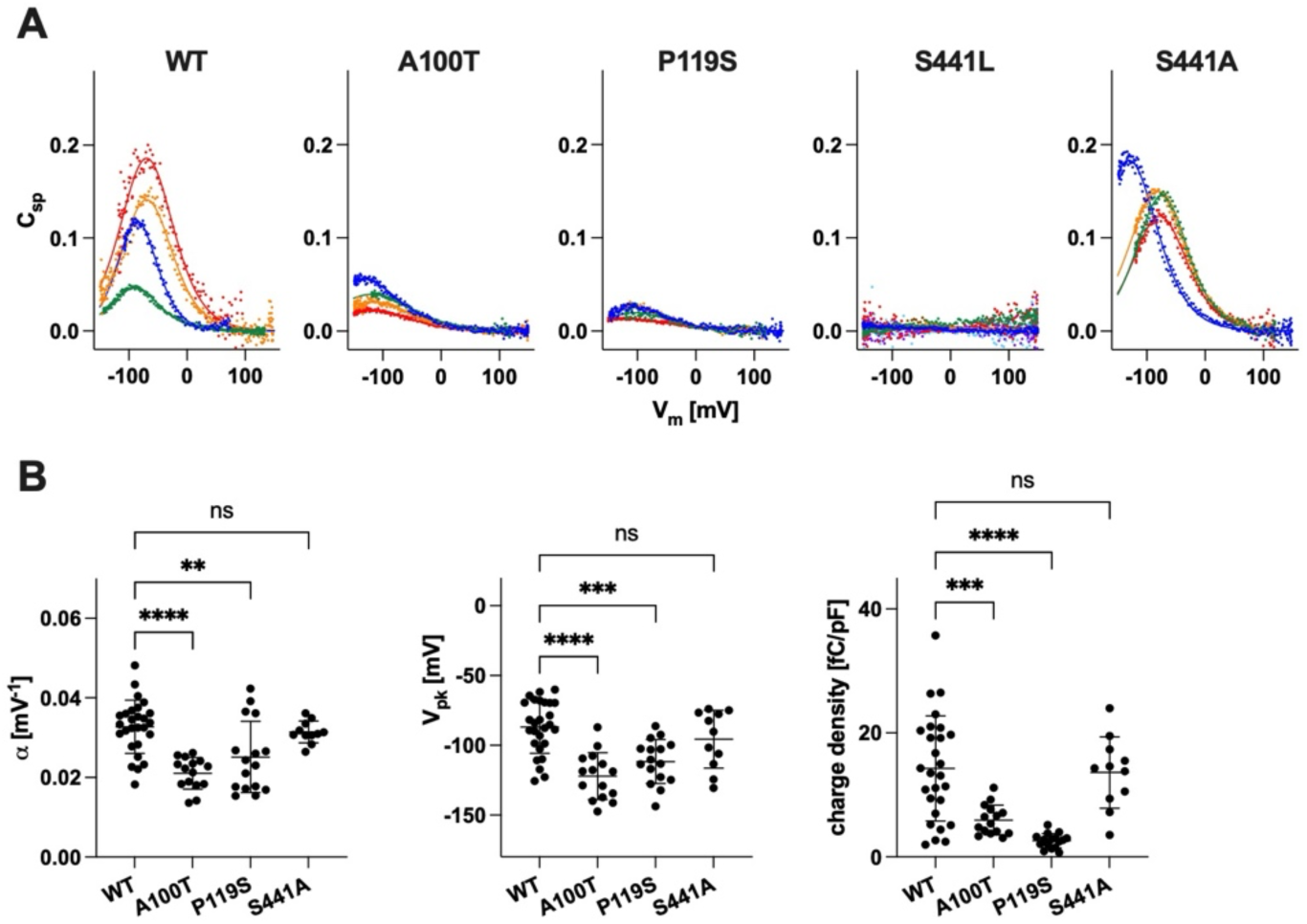
NLC measurements. (**A**) Examples of NLC recorded in HEK293T cells expressing WT, p.A100T, p.P119S, p.S441L, or p.S441A prestin. Different colors indicate individual recordings. (**B**) Summaries of the NLC parameters (α, V_pk_, and CD). Error bars indicate SD. ns, *p* ≥ 0.05; **, 0.001< *p* ≤ 0.01; ***, 0.0001< *p* ≤ 0.001; ****, *p* ≤ 0.0001.

**Fig. 5.**
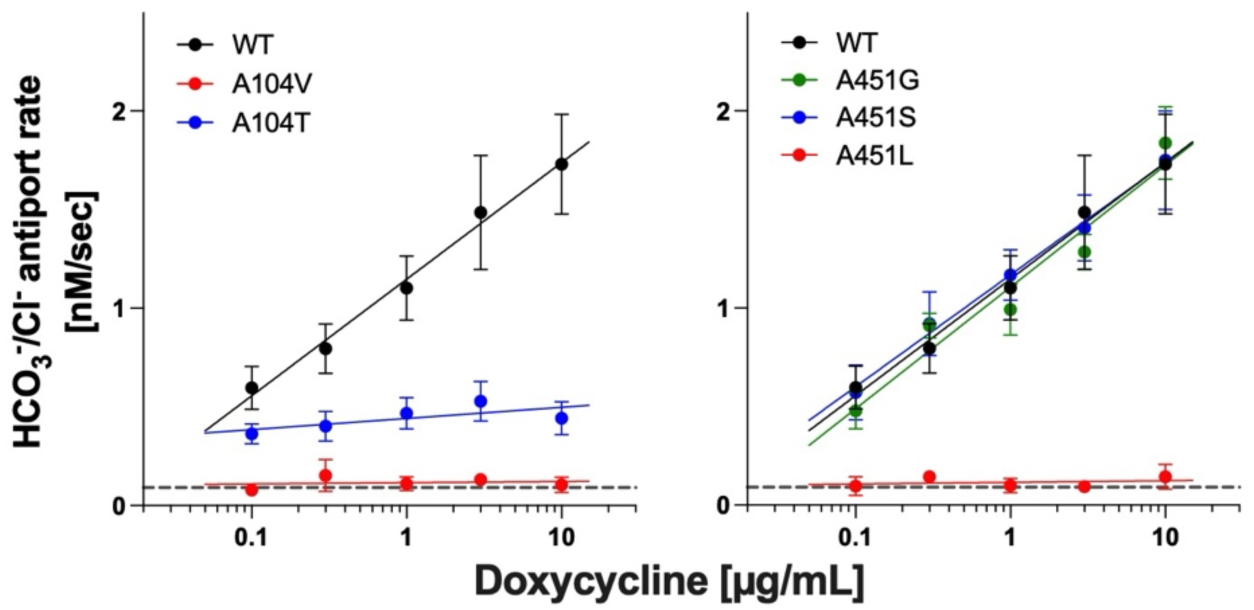
The effects of missense changes at Ala^104^ and Ala^451^ on the anion transport function of pendrin. HCO_3_^-^/Cl^-^ antiport assay conducted for p.A104V (left), p.A104T (left), p.A451G (right), p.A451S (right), and p.A451L (right) pendrin alongside WT control. Error bars indicate SD. Horizontal dotted lines indicate transport rates of uninduced cells. Solid lines indicate linear regressions (log_10_ [Dox] vs. transport rates). Sample size information and statistics are provided in Table 1.

Unlike p.A100T and p.P119S, NLC of p.S441L prestin was undetectable, indicating that p.S441L completely abrogates the motor function of prestin (**Figs. 4A** and **4B**). Ser^441^ is located at the interface between the gate and core domains (**Fig. 3A**, **right**). Equivalent residues at this site in the other SLC26 family members are either Ala or Gly (Ala^451^ in pendrin). These small residues may be important not to hinder the relative elevator-like movements of the core domain with respect to the gate domain. If true, p.S441L may sterically interfere with this interdomain motions. In line with this speculation, p.A451L in pendrin (equivalent to p.S441L in prestin) abolished the anion transport activity, whereas p.A451S and p.A451G did not (**Fig. 5, right**). We also found that p.S441A prestin has WT-like NLC (**Figs. 4A** and **4B**). These observations suggest the common importance of having a small residue at the Ser^441^ site in prestin and equivalent sites in the other SLC26 proteins.

### Comparison to AlphaMissense variant effect predictor

The pathogenicity of disease-associated variants is best assessed by functional analyses such as those presented here, however, experimental characterization of variants is laborious and remains reactive to the rapid identification of novel variants. Recent advancement in computational methods using machine learning has prompted us to evaluate the performance of AlphaMissense (AM), a novel tool for prediction of pathogenicity of missense variants (Cheng et al., 2023). First, we visualized the provided AM prediction scores of pendrin and prestin at every amino acid positions as heatmaps (**Supplementary Figs. 1** and **2**). AM pathogenicity predictions for both pendrin and prestin shows similarity, such as i) variants in regions that lack structural information (C- and N-termini, IDR) tends to be benign (i.e., predominantly “blue”); and ii) variants in transmembrane regions tend to be less tolerant to missense changes (i.e., predominantly “red”) in the heatmap (**Supplementary Figs. 1** and **2**). These observations are in line with the fact that AM utilizes structural information, and both pendrin and prestin shares quite similar overall molecular architecture. To evaluate the accuracy of AM, we plotted the HCO_3_^-^/Cl^-^ antiporter activities of pendrin variants measured in our previous study (Wasano et al., 2020) and this study, which are normalized to WT control, against the AM pathogenicity scores (**Fig. 6**). We found an overall correlation between our functional assay data and AM prediction scores of ρ = -0.6497 (nonparametric Spearman correlation), which is in line with the example given in their report (GCK, ρ = -0.65, Fig. 3G in Cheng *et al*.) (Cheng et al., 2023). Many variants tested in this study are clustered around highly conserved TM9-10 that are essential for the transport function, and thus many had little or no activities, skewing the distribution of variants (**Fig. 6**, **inset**). In addition, we found several variants that were mis-classified by AM as benign when there is vastly reduced activity (for example, p.N392S) or *vice versa* (ambiguous/pathogenic when their activity is WT-like, such as p.T307A, p.L117F, and p.F354S) (**Table 2**). For prestin variants, AM scores for p.A100T, p.P119S, and p.S441L were 0.9138, 0.7712, and 0.916, respectively, all predicting to be pathogenic.

**Fig. 6.**
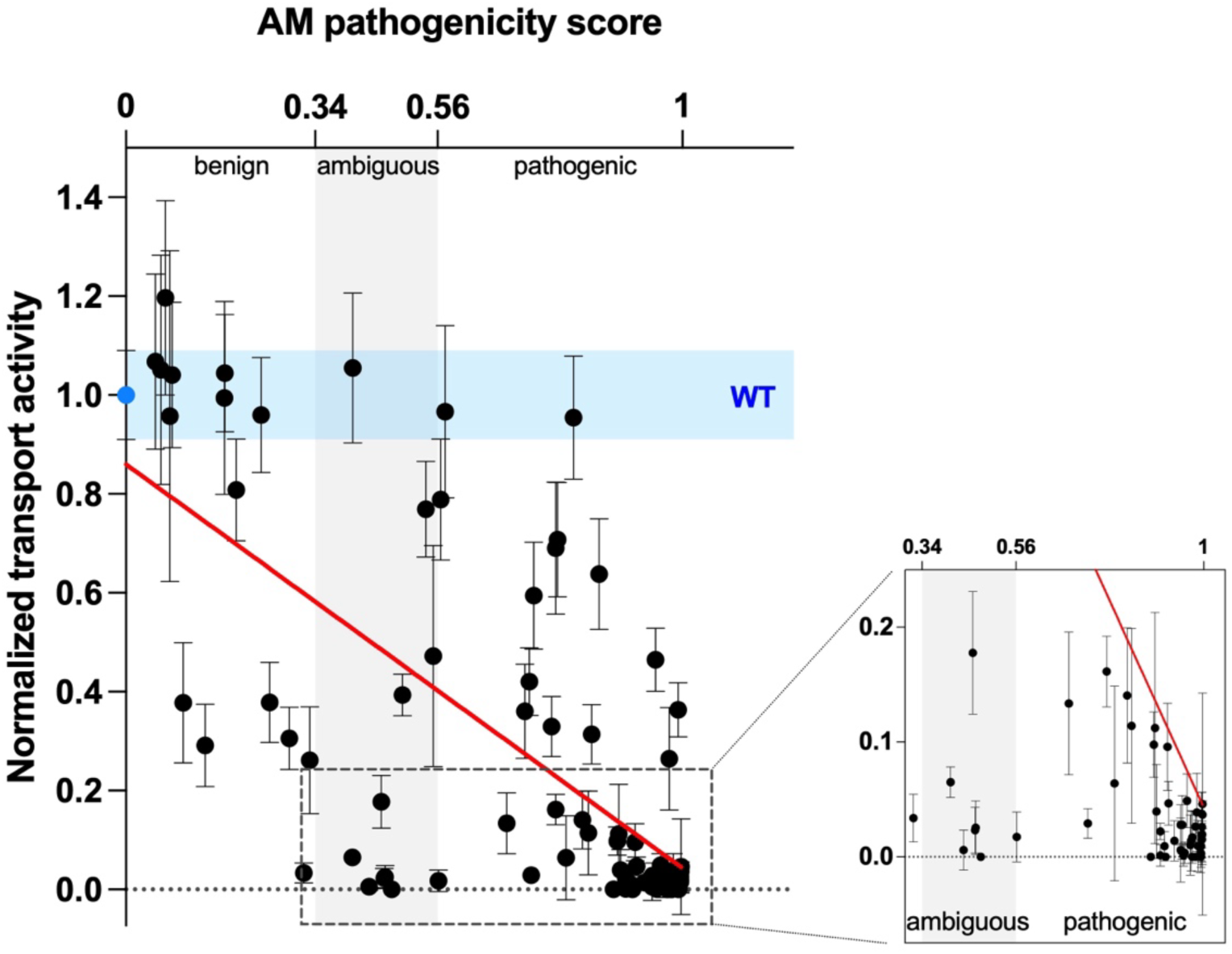
Correlation with AlphaMissense pathogenicity scores. HCO_3_^-^/Cl^-^ transport rates of pendrin WT and variants from this study and the previous study (Wasano et al., 2020) were normalized to the WT value and plotted against AlphaMissense (AM) pathogenicity scores (Cheng et al., 2023). Light blue shades indicate HCO_3_^-^/Cl^-^ rate of WT with errors (propagated errors). Gray shade between AM scores 0.34-0.56 marks the “ambiguous” category, with scores lower being “benign” and higher being “pathogenic” as indicated. Red line indicates the linear fit between the HCO_3_^-^/Cl^-^ antiport activity and the AM scores. Inset: Region indicated by the broken lines in left are enlarged to visualize variants with little or no HCO_3_^-^/Cl^-^ transport activity.

**Table 2.** Comparison of experimental data vs. AM scores. The slope values of the Dox-dependent HCO_3_^-^/Cl^-^antiport activities from our previous study (Wasano et al., 2020) and from this study for each variant were normalized to WT. For variants with negative slope values (slope < 0), normalized activity of 0 was assigned (highlighted in yellow) with no error propagations. Variants with WT-like HCO_3_^-^/Cl^-^ antiport activity (*p* ≥ 0.05) were shaded in light blue, and ones with impaired functions (*p* < 0.05) are shaded in pale pink. AM scores for pathogenicity classification uses the same color shades as in Supplementary Figs. 1 and 2. Asterisks (*) indicate variants not found in human patients (as of January 2024).

|  |  | HCO<sub>3</sub><sup>-</sup>/Cl<sup>-</sup> antiport assay results |  | AM prediction |  |
| --- | --- | --- | --- | --- | --- |
|  |  | %WT Activity | Variant Effect | AM Score | AM Category |
| Wasano et al., 2020 | p.Ser28Gly (c.82A>G) | 29.2 ± 8.3 | Function impaired | 0.1421 | benign |
|  | p.Ser49Arg (c.147C>G) | 104.1 ± 13.7 | WT-like | 0.0831 | benign |
|  | p.Pro76Ser (c.226C>T) | 30.5 ± 6.3 | Function impaired | 0.2937 | benign |
|  | p.Ser90Leu (c.269C>T) | 0.1 ± 0.9 | Function impaired | 0.8985 | pathogenic |
|  | p.Thr99Arg (c.296C>G) | 0 | Function impaired | 0.9802 | pathogenic |
|  | p.Leu117Phe (c.349C>T) | 95.5 ± 12.5 | WT-like | 0.804 | pathogenic |
|  | p.Pro123Ser (c.367C>T) | 2.9 ± 1.3 | Function impaired | 0.7286 | pathogenic |
|  | p.Gly131Val (c.392G>T) | 0.4 ± 1.7 | Function impaired | 0.9925 | pathogenic |
|  | p.Ser133Thr (c.397T>A) | 2.3 ± 2.0 | Function impaired | 0.4642 | ambiguous |
|  | p.Gly139Ala (c.416G>C) | 0 | Function impaired | 0.9105 | pathogenic |
|  | p.Met147Thr (c.440T>C) | 0.6 ± 2.8 | Function impaired | 0.9459 | pathogenic |
|  | p.Met147Val (c.439A>G) | 6.4 ± 8.5 | Function impaired | 0.7909 | pathogenic |
|  | p.Val163Ile (c.487G>A) | 105.1 ± 23.2 | WT-like | 0.0632 | benign |
|  | p.Val186Phe (c.556G>T) | 0 | Function impaired | 0.4777 | ambiguous |
|  | p.Thr193Ile (c.578C>T) | 0.9 ± 1.3 | Function impaired | 0.9082 | pathogenic |
|  | p.Tyr214Cys (c.641A>G) | 31.4 ± 6.0 | Function impaired | 0.8371 | pathogenic |
|  | p.Val239Asp (c.716T>A) | 4.9 ± 2.3 | Function impaired | 0.9612 | pathogenic |
|  | p.Asp266Asn (c.796G>A) | 106.7 ± 17.7 | WT-like | 0.0528 | ambiguous |
|  | p.Thr307Ala (c.919A>G) | 105.5 ± 15.2 | WT-like | 0.4074 | ambiguous |
|  | p.Asn324Tyr (c.970A>T) | 99.4 ± 19.5 | WT-like | 0.1773 | pathogenic |
|  | p.Gly334Val (c.1001G>T) | 59.5 ± 10.8 | Function impaired | 0.7329 | pathogenic |
|  | p.Phe354Ser (c.1061T>C) | 96.6 ± 17.4 | WT-like | 0.5739 | pathogenic |
|  | p.Lys369Glu (c.1105A>G) | 39.4 ± 4.2 | Function impaired | 0.4968 | ambiguous |
|  | p.Ala372Val (c.1115C>T) | 0.1 ± 0.9 | Function impaired | 0.9528 | pathogenic |
|  | p.Asn392Tyr (c.1174A>T) | 0.5 ± 0.7 | Function impaired | 0.9493 | pathogenic |
|  | p.Ser399Pro (c.1195T>C) | 78.9 ± 12.2 | Function impaired | 0.5658 | pathogenic |
|  | p.Ser408Phe (c.1223C>T) | 0.9 ± 1.3 | Function impaired | 0.9938 | pathogenic |
|  | p.Arg409His (c.1226G>A) | 2.8 ± 2.6 | Function impaired | 0.9499 | pathogenic |
|  | p.Thr410Lys (c.1229C>A) | 1.9 ± 1.1 | Function impaired | 0.9957 | pathogenic |
|  | p.Thr410Met (c.1229C>T) | 1.4 ± 1.7 | Function impaired | 0.9314 | pathogenic |
|  | p.Thr416Pro (c.1246A>C) | 2.2 ± 0.7 | Function impaired | 0.8985 | pathogenic |
|  | p.Gln421Leu (c.1262A>T) | 14.1 ± 5.9 | Function impaired | 0.8205 | pathogenic |
|  | p.Ile426Asn (c.1277T>A) | 63.8 ± 11.2 | Function impaired | 0.8501 | pathogenic |
|  | p.Leu445Trp (c.1334T>G) | 0 | Function impaired | 0.9933 | pathogenic |
|  | p.Asn457Lys (c.1371C>A) | 36.4 ± 5.5 | Function impaired | 0.992 | pathogenic |
|  | p.Arg470His (c.1409G>A) | 119.6 ± 19.7 | WT-like | 0.0709 | benign |
|  | p.Val483Glu (c.1448T>A) | 70.7 ± 11.6 | WT-like | 0.7757 | pathogenic |
|  | p.Gly497Ser (c.1489G>A) | 9.8 ± 2.9 | Function impaired | 0.883 | pathogenic |
|  | p.Thr527Pro (c.1579A>C) | 42.0 ± 6.8 | Function impaired | 0.7253 | pathogenic |
|  | p.Ile529Ser (c.1586T>G) | 16.1 ± 3.1 | Function impaired | 0.7726 | pathogenic |
|  | p.Tyr556Cys (c.1667A>G) | 6.5 ± 1.3 | Function impaired | 0.4069 | ambiguous |
|  | p.Cys565Tyr (c.1694G>A) | 80.8 ± 10.3 | WT-like | 0.1986 | benign |
|  | p.Ser657Asn (c.1970G>A) | 33.0 ± 6.0 | Function impaired | 0.7648 | pathogenic |
|  | p.Val659Leu (c.1975G>C) | 37.9 ± 8.1 | Function impaired | 0.2582 | benign |
|  | p.Ser666Phe (c.1997C>T) | 17.8 ± 5.3 | Function impaired | 0.4588 | ambiguous |
|  | p.Asp669Glu (c.2007C>A) | 0.0 ± 0.0 | Function impaired | 0.973 | pathogenic |
|  | p.Phe683Ser (c.2048T>C) | 2.8 ± 1.8 | Function impaired | 0.9461 | pathogenic |
|  | p.Phe692Leu (c.2074T>C) | 36.0 ± 9.6 | Function impaired | 0.7169 | pathogenic |
|  | p.Leu703Pro (c.2108T>C) | 0.3 ± 0.9 | Function impaired | 0.955 | pathogenic |
|  | p.Thr721Met (c.2162C>T) | 0.6 ± 1.7 | Function impaired | 0.4373 | ambiguous |
|  | p.His723Arg (c.2168A>G) | 13.4 ± 6.2 | Function impaired | 0.684 | pathogenic |
| This study | p.Gln383Glu (c.1147C>G) | 2.5 ± 2.4 | Function impaired | 0.4658 | ambiguous |
|  | p.Glu384Gly (c.1151G>C) | 2.7 ± 1.4 | Function impaired | 0.9808 | pathogenic |
|  | p.Ala387Val (c.1160C>T) | 4.7 ± 1.9 | Function impaired | 0.9179 | pathogenic |
|  | p.Gly389Arg (c.1165G>A) | 1.7 ± 1.5 | Function impaired | 0.996 | pathogenic |
|  | p.Gly389Arg (c.1165G>C) | 2.5 ± 3.1 | Function impaired | 0.996 | pathogenic |
|  | p.Ser391Asn (c.1172G>A) | 0 | Function impaired | 0.8762 | pathogenic |
|  | p.Ser391Arg (c.1173C>A) | 4.3 ± 2.4 | Function impaired | 0.9974 | pathogenic |
|  | p.Asn392Ser (c.1175A>G) | 3.4 ± 2.0 | Function impaired | 0.3199 | benign |
|  | p.Gly396Glu (c.1187G>A) | 1.0 ± 0.2 | Function impaired | 0.9866 | pathogenic |
|  | p.Val402Met (c.1204G>A) | 1.7 ± 2.2 | Function impaired | 0.5617 | pathogenic |
|  | p.Thr404Ile (c.1211C>T) | 1.7 ± 2.2 | Function impaired | 0.9734 | pathogenic |
|  | p.Ala406Thr (c.1216G>A) | 20.0 ± 4.9 | Function impaired | 0.8845 | pathogenic |
|  | p.Ser408Asp (c.1222_3TC>GA)* | 3.7 ± 1.2 | Function impaired | 0.9971 | pathogenic |
|  | p.Ser408Glu (c.1222_4TCC>GAG)* | 2.0 ± 2.0 | Function impaired | 0.9964 | pathogenic |
|  | p.Arg409Cys (c.1225C>T) | 3.4 ± 4.1 | Function impaired | 0.8893 | pathogenic |
|  | p.Arg409Leu (c.1226G>T) | 12.0 ± 2.8 | Function impaired | 0.9763 | pathogenic |
|  | p.Ala411Pro (c.1231G>C) | 0 | Function impaired | 0.9893 | pathogenic |
|  | p.Ala411Thr (c.1231G>A) | 77.0 ± 9.8 | WT-like | 0.5393 | ambiguous |
|  | p.Val412Ile (c.1234G>A) | 95.8 ± 33.5 | WT-like | 0.0792 | benign |
|  | p.Gln413Pro (c.1238A>C) | 3.9 ± 3.4 | Function impaired | 0.9839 | pathogenic |
|  | p.Gln413Arg (c.1238A>G) | 11.2 ± 10.0 | Function impaired | 0.8859 | pathogenic |
|  | p.Glu414Lys (c.1240G>A) | 11.3 ± 8.5 | Function impaired | 0.8309 | pathogenic |
|  | p.Ser415Gly (c.1243A>G) | 37.9 ± 12.1 | Function impaired | 0.1027 | benign |
|  | p.Ser415Arg (c.1245C>A) | 46.5 ± 6.4 | Function impaired | 0.9514 | pathogenic |
|  | p.Ser532Ile (c.1595G>T) | 26.2 ± 10.8 | Function impaired | 0.3304 | benign |
|  | p.Ala104Val (c.311C>T) | 1.0 ± 2.5 | Function impaired | 0.9684 | pathogenic |
|  | p.Ala104Thr (c.310G>A)* | 9.6 ± 3.8 | Function impaired | 0.9153 | pathogenic |
|  | p.Ala451Gly (c.1352C>G)* | 104.6 ± 11.8 | WT-like | 0.1774 | benign |
|  | p.Ala451Ser (c.1351G>T)* | 96.1 ± 11.6 | WT-like | 0.2429 | benign |
|  | p.Ala451Leu (c.1351_2GC>CT)* | 1.4 ± 2.5 | Function impaired | 0.9707 | pathogenic |

## Discussion

Precisely understanding the relationships between genotypes and disease phenotypes is the holy grail of the field of human genetics. Although the sequencing information becomes more and more accessible, assigning pathogenicity to variations of sequences remains challenging due to the complexity in determining the causality of variants to disease states (Brandes et al., 2022). Although experimental interrogation is ideal, our collective experimental efforts still have only scratched the surface on building an atlas of variant effects in the genome.

Consequently, the vast majority of disease-associated variants, including those of pendrin and prestin, remain as variants of unknown significance (VUS) (Richards et al., 2015). To bridge the gap, *in silico* tools have been developed to predict whether a variant plays a causative role in a disease state. Large numbers of variant effect predictors (VEPs) have been reported over the years using various approaches with various degrees of performances (Livesey and Marsh, 2022). For the deafness genes, a measure for protein-folding stability was used to re-classify VUSs in patients due to protein misfolding (Tollefson et al., 2023). Recently developed AlphaMissense (AM) also uses structural information, although it does not predict protein stability. Rather, it classifies missense variants into benign, ambiguous, or pathogenic based on their prediction scores (Cheng et al., 2023). As the study provided the AM prediction across the entire human proteome, we compared the available AM prediction scores for pendrin to our experimental results on quantitated transport activities. We found good correlation (ρ = -0.6497, **Fig. 6**), demonstrating their utility in informing the variant effects. It must be noted, however, that the uncertainty still exists as several pendrin variants were mis-classified. Thus, the usage of VEPs, even the one demonstrated superior performance over many existing VEPs across multiple benchmarks as AM, must exercise utmost caution in the clinical settings.

Recent efforts in large scale systemic studies (multiplexed assay of variant effects, MAVE) have reported functional consequences of hundreds of thousands of variants across genome regions to date (Fowler et al., 2023; Tabet et al., 2022). For proteins such as pendrin and prestin, however, large-scale multiplexing assays will be challenging as their dysfunctions may not result in easily discernable cellular phenotypes. Nevertheless, our small-scale *in vitro* assays are robust in determining the degrees of functional impairment of the variants found in human patients. The large variation in PDS and DFNB4 phenotypes suggests large variations in activities of pendrin variants, thus warranting the experimental approach that can detect a subtle difference in transport activities. One caveat will be that our approach in heterologous expression system may not be able to address effects of variants on protein expression levels in the host environment, as we forcibly induce protein expression by doxycycline from cDNAs for our transporter assays. Also not addressed here is the variant effects on protein-protein interactions. For example, the intrinsically disordered region (IDR) within the cytosolic domain in prestin mediate binding of calmodulin thereby allowing modulation of protein activity by calcium, which is likely conserved in other SLC26 proteins including pendrin (Keller et al., 2014). Additionally, IQ-motif containing GTPase-activating protein 1 (IQGAP1) was shown to interact with the C-terminal region of pendrin and enhances its transport activity in kidney cells (Xu et al., 2022). Interestingly, co-expression of IQGAP1 and pendrin also enhanced the plasma membrane (PM) localization of pendrin, suggesting another layer of regulatory interaction. It is curious that although both pendrin and prestin are conserved and the handful of disease-associated variants in prestin also similarly affected pendrin in this study, prestin is targeted to the lateral membrane in the outer hair cells (OHCs) while pendrin is targeted to the apical membranes of epithelial cells in thyroid and in the inner ear (Royaux et al., 2000; Wangemann et al., 2004; Zheng et al., 2005; Zheng et al., 2010). Thus, it will require additional considerations such as co-expression of interacting partners or using the host cell lines in order to interrogate the effects of variants found in regions where such regulatory interactions were reported in the future.

Although functional annotation of deafness-associated variations in the genome may not be complete until the hearing phenotype is assessed *in vivo*, our continuing efforts in characterizing missense variants *in vitro* provide invaluable information regarding the variant effects, which can be used to prioritize future efforts at the organismal level. With ever improving VEPs such as AM, future studies will provide clinically useful information of variant effects with more confidence.

## ABBREVIATIONS

SLC: solute carrier;
PDS: pendred syndrome;
EVA: enlarged vestibular aqueduct;
OHC: outer hair cell;
NLC: nonlinear capacitance;
TM: transmembrane;
WT: wild type;
mTq2: mTurquoise2;
VUS: variants of unknown significance;
VEP: variant effect predictor;
AM: AlphaMissense

## Acknowledgement

This study is supported by NIH grant DC017482 to K. H. and the Hugh Knowles Center.

## Author Contributions

K.H. conceived the study; S.T., T.K., K.W., and K.H. performed experiments; S.T., and K.H. analyzed the data; S.T., and K.H. wrote the manuscript.

---

**Supplemental Fig. 1.**
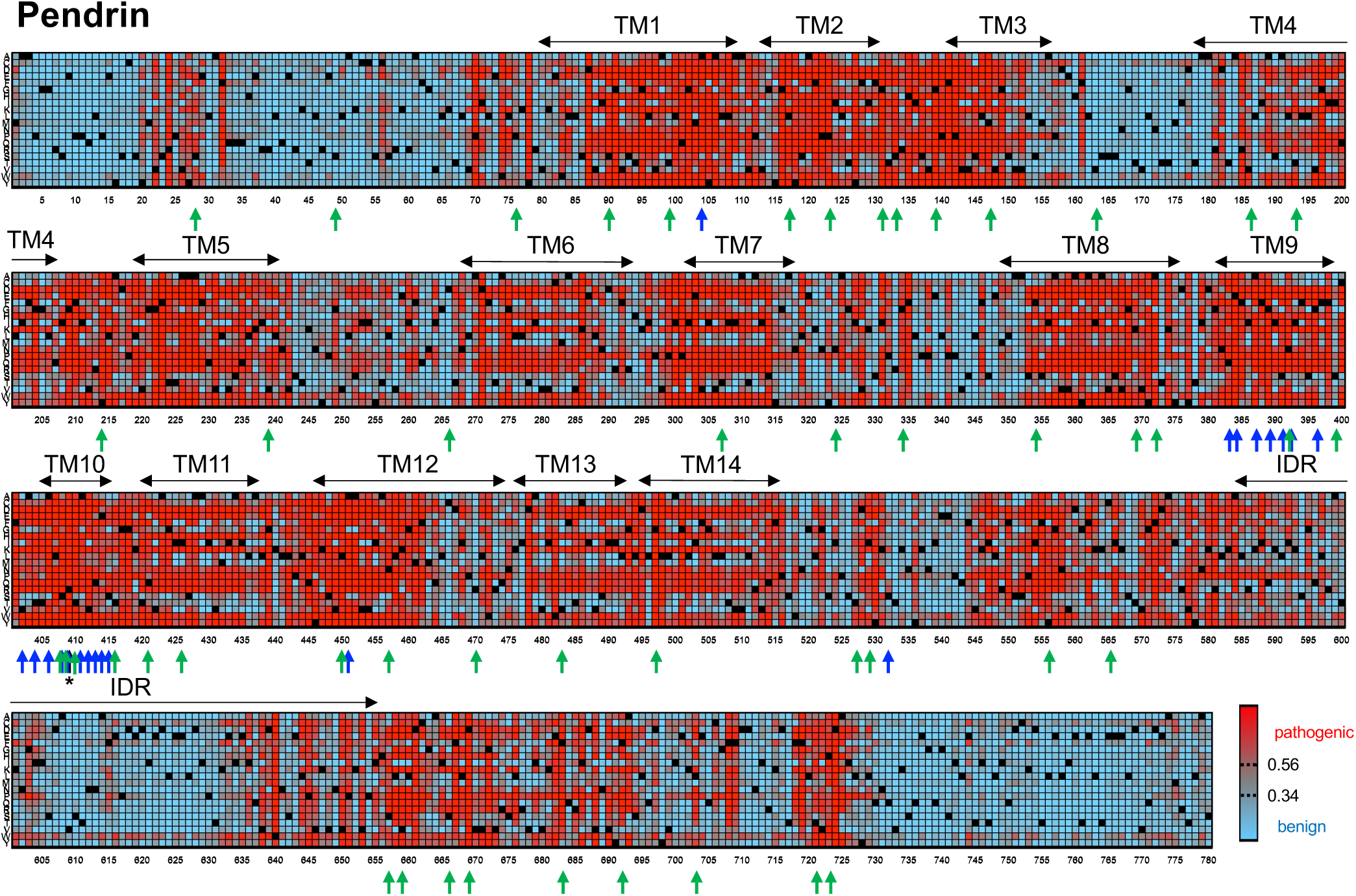
AlphaMissense pathogenicity scores of pendrin. Heatmap showing the AM scores for the entire length of human pendrin. Variants with a score from 0-0.34 are classified as benign (cyan), 0.34-0.56 as ambiguous (gray), and 0.56-1.0 are pathogenic (red) by AM. Amino acid residue numbers are shown on the bottom and the amino acids (one letter code) are indicated on the left. WT residues are shown in black. Upward arrows indicate the positions of the residues with the variants tested in the previous study (green, (Wasano et al., 2020) or in this study (blue). Black upward arrow with an asterisk indicates the anion binding site. Horizontal double arrows show the regions that comprise the transmembrane (TM) helices and the intrinsically disordered region (IDR).

**Supplemental Fig. 2.**
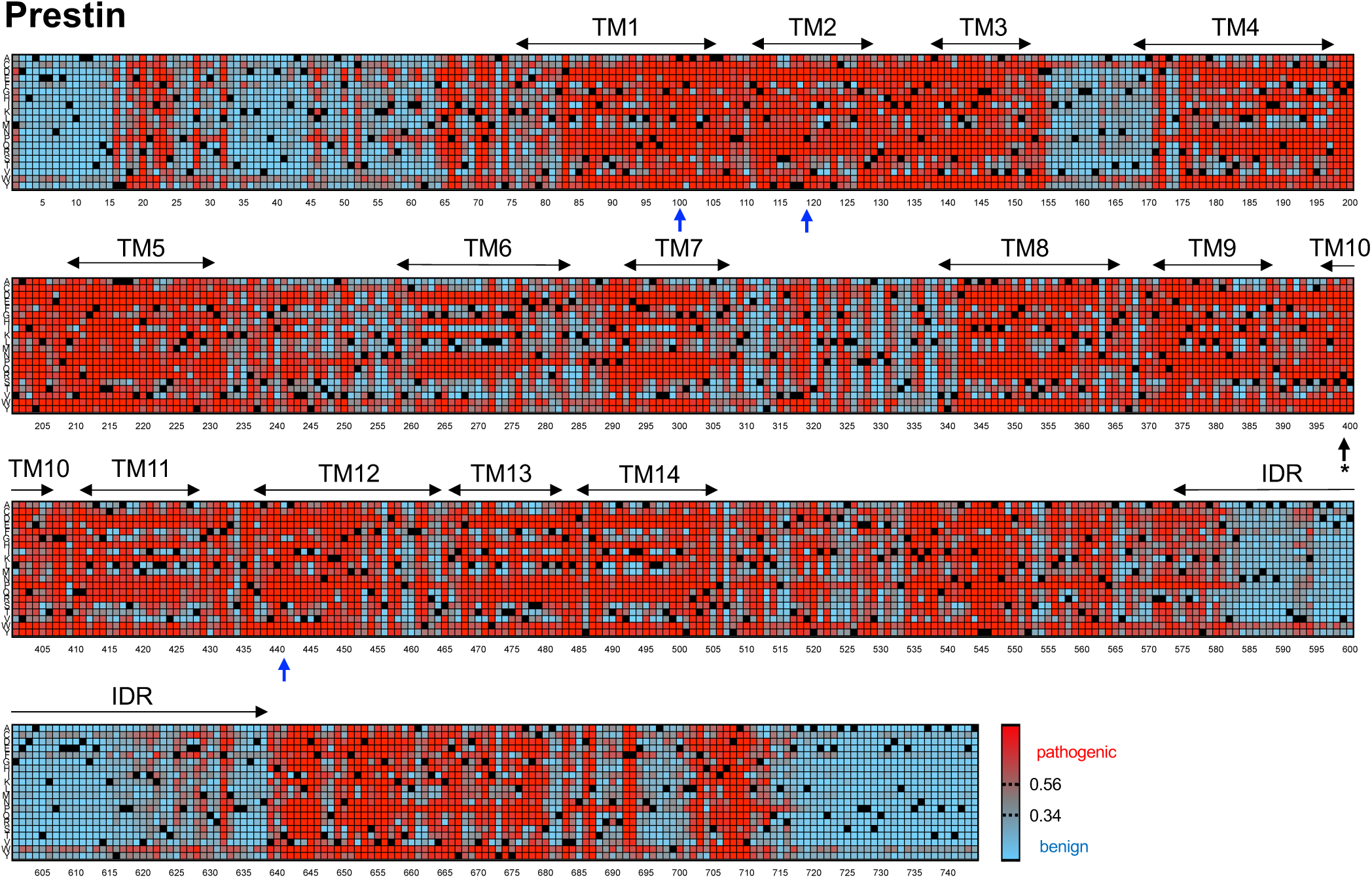
AlphaMissense pathogenicity scores of prestin. Heatmap showing the AM scores for the entire length of human prestin. AM pathogenicity scores are colored as in Supplemental Fig. 1. Amino acid residue numbers are shown on the bottom and the amino acids (one letter code) are indicated on the left. WT residues are shown in black. Blue upward arrows indicate the positions of the residues with variants tested in this study. Black upward arrow with an asterisk indicates the anion binding site. Horizontal double arrows show the regions that comprise the transmembrane (TM) helices and the intrinsically disordered region (IDR).

